# BDdb: a comprehensive platform for exploration and utilization of birth defect multi-omics data

**DOI:** 10.1101/2020.07.20.211391

**Authors:** Dengwei Zhang, Hai-Xi Sun, Ziheng Zhou, Xiaosen Jiang, Dongsheng Chen, Si Zhou, Jie Huang, Shoufang Qu, Ying Gu, Xiuqing Zhang, Xin Jin, Ya Gao, Yue Shen, Fang Chen

**Affiliations:** BGI-Shenzhen, Shenzhen 518083, China; Guangdong Provincial Key Laboratory of Genome Read and Write, BGI-Shenzhen, Shenzhen, 518120, China; Shenzhen Institute of Synthetic Biology, Shenzhen Institutes of Advanced Technology, Chinese Academy of Sciences, Shenzhen, People’s Republic of China; China National Genebank, BGI-Shenzhen, Shenzhen 518120, China; BGI Education Center, University of Chinese Academy of Sciences, Shenzhen 518083, China; BGI Genomics, BGI-Shenzhen, Shenzhen 518083, China; National Institutes for food and drug Control (NIFDC), Beijing 100050, P. R. China; Shenzhen Engineering Laboratory for Innovative Molecular Diagnostics, BGI-Shenzhen, Shenzhen, 518120, China; Shenzhen Engineering Laboratory for Birth efects Screening, Shenzhen, 518120, China; Guangdong Provincial Key Laboratory of Human Disease Genomics, Shenzhen Key Laboratory of Genomics, BGI-Shenzhen, Shenzhen 518083, China; Shenzhen Engineering Laboratory for Birth Defects Screening, BGI-Shenzhen, Shenzhen 518083, China

**Author notes:** Contribute equally to this work (Dengwei Zhang, Hai-Xi Sun, Ziheng Zhou, Xiaosen Jiang, Dongsheng Chen). Corresponding authors Correspondence should be addressed to Fangchen, Yue Shen, Ya Gao, Xin Jin.

## Abstract

Birth defect, not only poses a major challenge for infant health but also attracts the attention of countless people in the world. Chromosome abnormality directly results in diverse birth defects which are generally deleterious and even lethal. Therefore, gaining molecular regulatory insights into these diseases is important and necessary for effective prenatal screening. Recently, with the advance of next-generation sequencing (NGS) techniques, a myriad of treatises and data associated with these diseases are now constantly produced from different laboratories across the world. To meet the increasing requirements for birth-related data resources, we developed a birth defect multi-omics database (BDdb), freely accessible at http://t21omics.cngb.org and consisting of multi-omics data, circulating free DNA (cfDNA) data, as well as diseases biomarkers. Omics data sets from 138 GSE samples, 5271 GSM samples and 328 entries, and more than 2000 biomarkers of 22 birth-defect diseases in 5 different species were integrated into BDdb, which provides a user-friendly interface for searching, browsing and downloading selected data. Additionally, we re-analyzed and normalized the raw data so that users can also customize the analysis using the data generated from different sources or different High-Throughput Sequencing (HTS) methods. To our knowledge, BDdb is the first comprehensive database associated with birth-defect-related diseases. which would benefit the diagnosis and prevention of birth defects.

## INTRODUCTION

Birth defects (BD) refer to abnormalities in form, function, biochemistry, and mentality that exist after the baby leaves the mother’s body. More than 8.14 million children are born with severe BD each year, and BD has become one of the main causes of infant death and congenital disability in the world (1–3). The impacts of BD on human society are widespread: it not only seriously affects children’s survival and quality of life, but also affects the happiness and harmony of the family, and even causes a socioeconomic burden. Although there have been many studies aiming to understand the relationship between various risk factors and the incidence of BD, ~60% of the cause of BD in human still remains unclear. The known causes of BD can be roughly divided into three categories: genetic factors, environmental factors, and the combined effects of genetic and environmental factors. Among them, genetic factors are the predominant cause, which include BD caused by genetic mutations or chromosomal aberrations, for example, an increase or decrease in the number of chromosomes, structural abnormalities, and the duplication and loss of genetic material leading to severe mental retardation or deformity (3,4).

Chromosomal abnormalities are one of the main causes of BD. It is estimated that aneuploid chromosomal diseases account for about 6% of BD, and nearly 1 in 200 newborns are affected (5). Chromosome aneuploidy is defined as an increase or decrease in chromosomes within an individual that is not a multiple of the haploid. Most of these patients suffer severe intellectual disability and deformities of tissues and organs. The most common chromosome abnormalities include trisomy 21 (Down’s syndrome), trisomy 18 (Edward syndrome), trisomy 13 (Patau syndrome), X monomer (45, X, Turner syndrome), 47, XXY (Klinefelter syndrome), triploid and chimera, etc., conferring more than 80% of prenatal diagnosis of chromosomal abnormalities. Among them, Down syndrome is the most common disease whose incidence is 1/800-1/600 in the population. There are many types of diseases involved in BD with at least 8,000-10,000 are currently known. Because almost no effective treatment is available, it has become a public health issue that affects the quality of the population and the health of the group (6).

So far, chromosome diseases have almost no effective treatment. Only early diagnostic strategies, such as prenatal screening and prenatal diagnosis, can be performed during pregnancy to prevent BD. Therefore, understanding the molecular mechanisms of diseases associated with BD and the function of pathogenic genes is extremely important for the treatment of these diseases and the related research. However, there is currently no multi-omics database for collating, storing, integrating, and displaying BD-related research data sets. To shed light on this, we build BDdb, a user-friendly database that not only allows users to effectively view each date set, but also to mine deeper information through comparison of multiple data sets.

The current version of BDdb includes more than 100 manually curated datasets in human, involving 36 cell types and 13 tissues, as well as 12 tissues and 17 cell lines in mouse. This database provides a powerful platform that allows for quick retrieval of datasets of particular cell types in any tissue of interest. The main functions of the database include search, displaying differential gene lists, GO and KEGG enrichment analysis, genome browser, protein interaction network, etc. Furthermore, BDdb deposits more than 2000 manually curated biomarkers in 6 species, involving 22 birth-defect diseases. In summary, BDdb provides a comprehensive resource of BD for researchers and clinicians conveniently.

## MATERIALS AND METHODS

### Data collection

Since chromosome abnormality bears relation to the birth defect greatly, we firstly collected the data of this field. The data sets used to build BDdb were collected from Gene Expression Omnibus (GEO, https://www.ncbi.nlm.nih.gov/geo/) database, and we employed a list of keywords such as “trisomy 21”, “trisomy 13”, “trisomy 18”, “trisomy 8”, “monosomy X” and “XXY” to retrieve pertinent information in *Homo sapiens* and *Mus musculus*, by which we obtained 328 GSE samples in total (Figure 1). All datasets were collected before August 2019. We then manually screened out the relevant GSE samples with a careful reading of each retrieval data. Eventually, a total of 138 GSE samples were picked out and re-analyzed.

**Figure 1.**
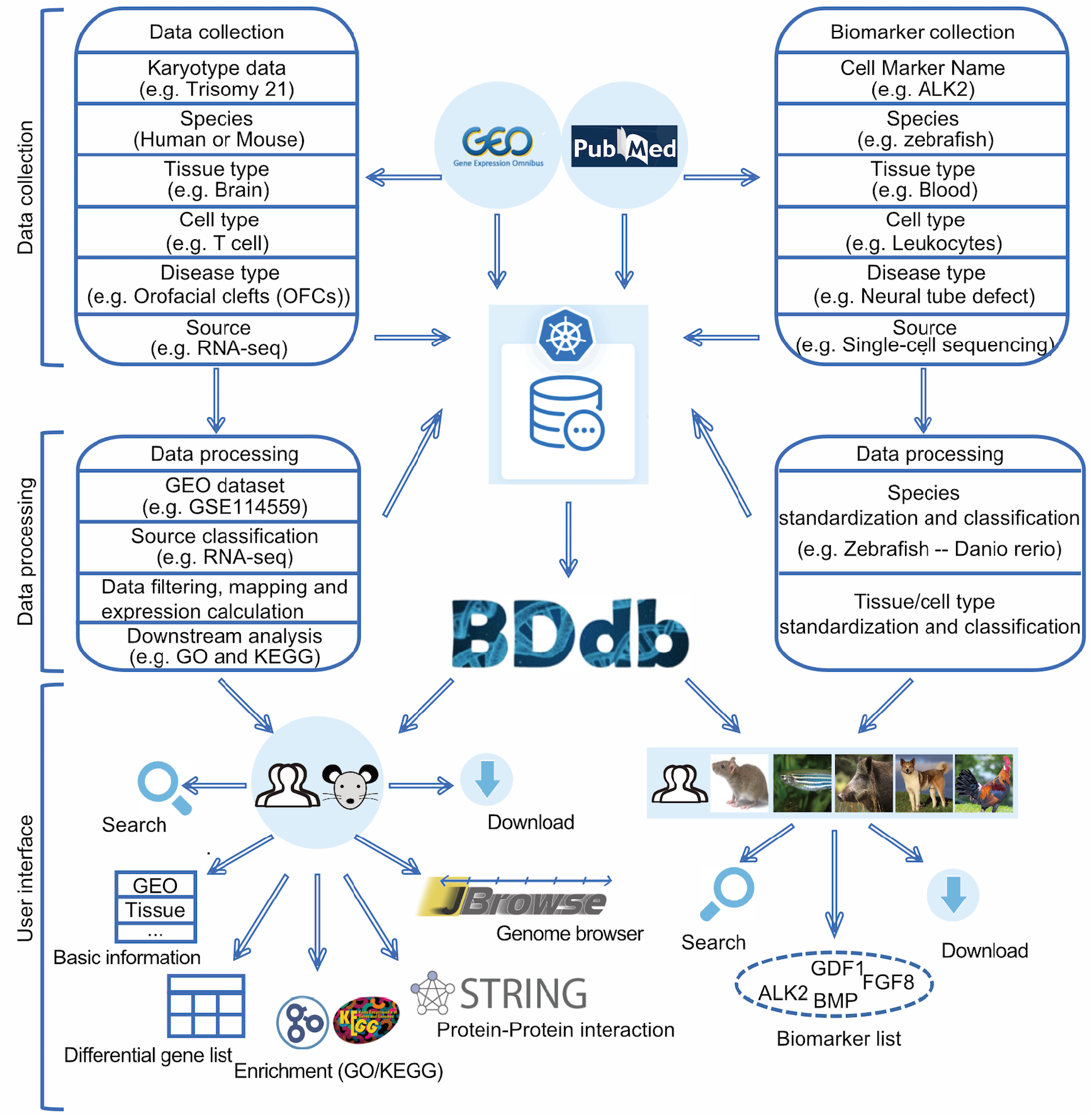
Overview of data collection, processing and database interface.

In 1997, fetal cell-free DNA was found in maternal peripheral blood plasma (1), opened up a variety of new possibilities for non-invasive prenatal testing capability. For example, maternal plasma DNA can be used as a safe, accurate and rapid non-invasive detection to determine whether the fetus has Down’s syndrome or other birth defects (2). We searched NCBI by a series of keywords, including ‘cell-free DNA’ and ‘cell-free RNA’, and collected more than 1000 datasets and related publications. Then, we carefully read the summary of these datasets and the introduction of each paper, and then we identified the datasets related to birth-defect, which were used in the subsequent analysis.

To ensure comprehensive data collection, we searched NCBI using keywords, such as ‘peripheral whole blood’, ‘PBMC’ and ‘peripheral blood’. We filtered out the datasets that coincide with the first collection and ensured the retained data related to birth defects, and then we manually summed up related datasets and downloaded raw data for subsequent analysis.

Considering single-cell RNA sequencing (scRNA-seq) technologies opening a hot field that holds tremendous potential to study cell phenotype and cell behavior at single-cell resolution (7,8), we paid attention to the single-cell sequencing studies focusing on birth defect. We searched birth-disease related scRNA-seq datasets in human and mouse from Sequence Read Archive (SRA) and Gene Expression Omnibus (GEO), and we found one dataset (GSE127257) (9) including 13,766 cells. This dataset focused on Ts65Dn mouse model for DS, which has triplication of sequences syntenic with human chr21.

Furthermore, to obtain biomarkers associated with different birth-defect diseases from publications, we searched PubMed by keywords such as particular disease names with “marker(s)”, “gene(s)” and “genetic(s)”. A total of more than 3000 publications were obtained. We manually surveyed these publications’ abstract, and if studies were identified to include pertinent information, we would download the paper as well as its supplementary materials. Then, we further manually checked the full texts of these selected publications to collect biomarkers with reliable evidence and gained the useful information including the title of paper, PMID, disease type, sequencing type, tissue type, cell type and species. Among the final results, more than 500 publications relative to birth-defect disease marker genes were separately conducted and marker gene lists were manually extracted.

### Data processing

Raw data of next-generation sequencing and microarray were downloaded and re-analyzed, using a uniform pipeline to reduce and eliminate the variation caused by different analysis approaches among published papers. For the GSE samples without raw data, their original tables attached in GEO were used.

For each dataset, we carefully read the original paper if it is available and focused on the Method part of the paper. These “experimental design” could help users to explore these datasets in different patients and karyotypes or models. When a dataset contained different karyotypes or models (or different types of sources, tissue and cell lines), we manually divided it into multiple sub-datasets (Figure 1). Then 328 (sub)-datasets were generated.

Some datasets had difficulties in re-analysis and/or data type messy problems, to overcome this we used GEO2R to obtain differentially expressed genes (DEGs) list, and then, we processed functional enrichment analysis using KEGG and GO. Finally, we constructed the putative regulatory pathways by correlation analysis (10).

#### RNA-Seq

Raw reads were filtered by SOAPnuke (11) with the parameters “-l 15 -q 0.2 -n 0.05”. Clean reads were then aligned to human (GRCh38.p12, downloaded from GENCODE) or mouse (GRCm38.p6, downloaded from GENCODE) reference genome using HISAT2 (12), and StringTie (13) was utilized to compute gene expression. To screen differentially expressed genes (DEGs), DESeq2 (14) was adopted when the sample had replicates; otherwise, DEGs were fetched using edgeR (15) if there were no replicates in the sample. Meanwhile, they were based on the screening condition that the absolute log_2_-fold change >= 0.5, and p-value < 0.05 and FDR < 0.1. Gene ontology (GO) enrichment and KEGG annotation were performed using clusterProfiler (16).

#### DNA methylation

The raw reads from BS-Seq or RRBS-Seq were filtered with Trim Galore (17) to trim low-quality bases and adapter. Subsequently, reads alignment and the extraction of methylation information were performed using Bismark (18). In particular, PCR duplications have been removed for BS-Seq sample, while RRBS-Seq sample skipped it in accordance with the Bismark’s tutorial. Furthermore, methylKit (19) was used to perform analyzing the differential methylation region among diverse samples.

#### DNA-protein interactions

For the analysis of ChIP-Seq and DNase-Seq, an identical pipeline was adopted. SOAPnuke (11) was employed to filter out low-quality reads, and clean reads were then mapped against human or mouse reference using Bowtie2 (20). Peak calling was performed by MACS2 (21,22) and peak annotation was accomplished using CHIPseeker (23). At last, DiffBind (24) was used to find out the differential peaks.

#### Small RNA (smRNA) Sequencing

Trim Galore (17) was adopted to trim adaptor of raw reads, and only the reads from smRNA ranging from 18 to 30 nt in length were kept with the parameters “--quality 20 --gzip --small_rna --max_length 30”. The eligible reads were then aligned against human reference genome (GRCh38.p12, downloaded from GENCODE) and the miRbase (25) (Release 22.1, http://www.mirbase.org/) to be annotated as miRNA, tRNA, rRNA, snoRNA, snRNA, piRNA or other non-coding RNAs.

#### Microarray

For DNA microarrays datasets with raw signal files, an in-house bioinformatics pipeline was adopted for quality control and expression quantification. Briefly, we read *cel* file by using R package affy (version 1.8 release) (26). Then, we used R package simpleaffy (27) and affyPLM (28) to assess chips quality, we filtered low-quality chips (actin3/actin5 >3, gapdh3/gapdh5 > 1.25 and non-detected BioB) and probe site with *lowvar* and feature exclude and then we use method ‘rma’ to process pretreatment. The expression abundance values of genes (including PCGs and lncRNAs) were summarized by using t-test. Expression values were log_2_ transformed with an offset of 1. We sorted out the list of up- and down-regulated genes, and then, we processed functional enrichment analysis using KEGG and GO. Finally, we constructed the network analysis of differentially expressed genes by STRING (10).

#### sci-ATAC-seq

We extracted the corresponding metadata, including the cell types and sources (i.e., tissue, cell line). Also, we obtained the cell groups (referring to patient ID, culture condition or cell phenotype) from the supplementary tables of the paper, and labeled each cell with the information of cell groups, and meanwhile, we collected the marker genes to help users to explore the birth defect disease using sci-ATAC-seq filed.

### Web construction

The web was constructed based on CodeIgniter, a powerful PHP framework. CodeIgniter provided an Application Programming Interface (API) to connect the web to MySQL database. We also used JavaScript libraries including jQuery (3.4.1), jQuery-labelauty and some additional plugins to perform dynamic web services.

### Data access

Now the BDdb website is accessible to users at http://t21omics.cngb.org, which offers a concise, well-organized interface. Meanwhile, users are welcomed to add comments regarding their requirements and suggestions in order to improve BDdb. All provided data can be downloaded. The BDdb will be updated continually and store more data from pertinent researches. Figure 1 shows the overall features of BDdb.

## RESULTS

### BDdb statistics

In total, BDdb contains 102 and 36 GSEs in human and mouse, respectively. Generally, data sets of 17 diseases, 13 tissues and 36 cell lines in human, as well as 4 diseases, 12 tissues and 17 cell lines in mouse, were included in BDdb (Figure 2A-E). These datasets are derived from 17 types of omics, including genomics, transcriptomics and epigenomics (Figure 2C, D). We also collected single-cell omics as well as cell-free DNA and RNA data sets, both of which are rapidly developing fields and have significant research values for birth defects. Notably, 66% and 83% of the data are derived from T21 in human and T65dn in mouse, which is the most karyotypic abnormality in this database. Moreover, BDdb supplements more than 2000 biomarkers belonging to 22 types of birth defects, including microcephaly, neural tube defect, preeclampsia, atrioventricular septal defect, and so on. These markers are derived from five species, including *Homo sapiens*, *Danio rerio*, *Mus musculus*, *Sus scrofa*, *Canis familiaris*, and *Gallus gallus* are obtained from more than 500 studies (Figure 2F).

**Figure 2.**
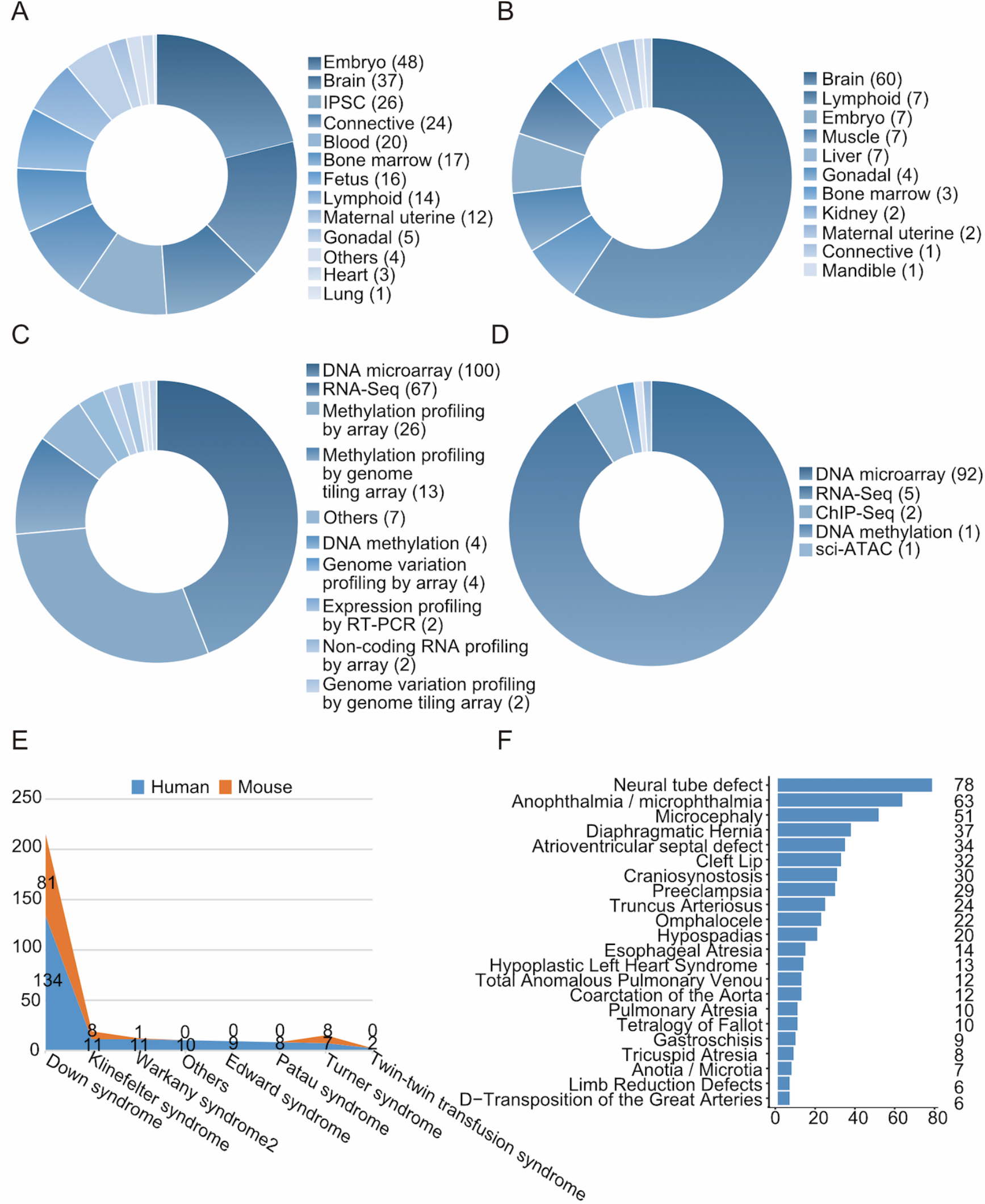
Statistics of datasets in BDdb. (A) Distribution of datasets for different tissue types in human. (B) Distribution of datasets for different tissue types in mouse. (C) Distribution of datasets for different omics in human. (D) Distribution of datasets for different omics in mice. (E) Distribution of datasets for different diseases in human and mouse. (F) The numbers of publications for different diseases.

BDdb is consisting of multi-omics datasets involving tens of common birth-defect diseases. Embryo tissue is the one with the most abundant samples in human, followed by brain and blood. Meanwhile, the brain takes the largest proportion in mouse tissues, whereas lymphoid and embryo place 2nd and 3rd, respectively (Figure 2A, B). Correspondingly, the most abundant cell lines are blastocysts belonging to embryo tissue in human and cortex belonging to brain tissue in mouse. For the diversity of sequencing types, DNA microarray tops in both human (44%) and mouse (91%), and the subsequent four are RNA-Seq, methylation profiling by array, ChIP-Seq and DNA methylation (Figure 2C, D). Moreover, the vast majority are linked to Down syndrome, followed by Klinefelter syndrome, Turner syndrome, Warkany syndrome 2 and Edward syndrome (Figure 2E). Additionally, BDdb reserves some data from birth-defect diseases not having associations with chromosome abnormalities, such as twin-twin transfusion syndrome, orofacial clefts (OFCs) and open myelomeningocele. For biomarkers collection, the top 5 diseases with most published studies are neural tube defect, anophthalmia/microphthalmia, microcephaly, diaphragmatic hernia, and atrioventricular septal defect (Figure 2F). A summary of these datasets and biomarkers can be found in Table 1 and Table 2, respectively.

### Database features and utility

BDdb collects birth-defect multi-omics data sets and allows users to query the subsequent analysis results with 5 functional states. The easy-to-use interface provides access for searching, browsing, visualizing and downloading (Figure 3). The online user guide illustrates several cases of BDdb usage.

**Figure 3.**
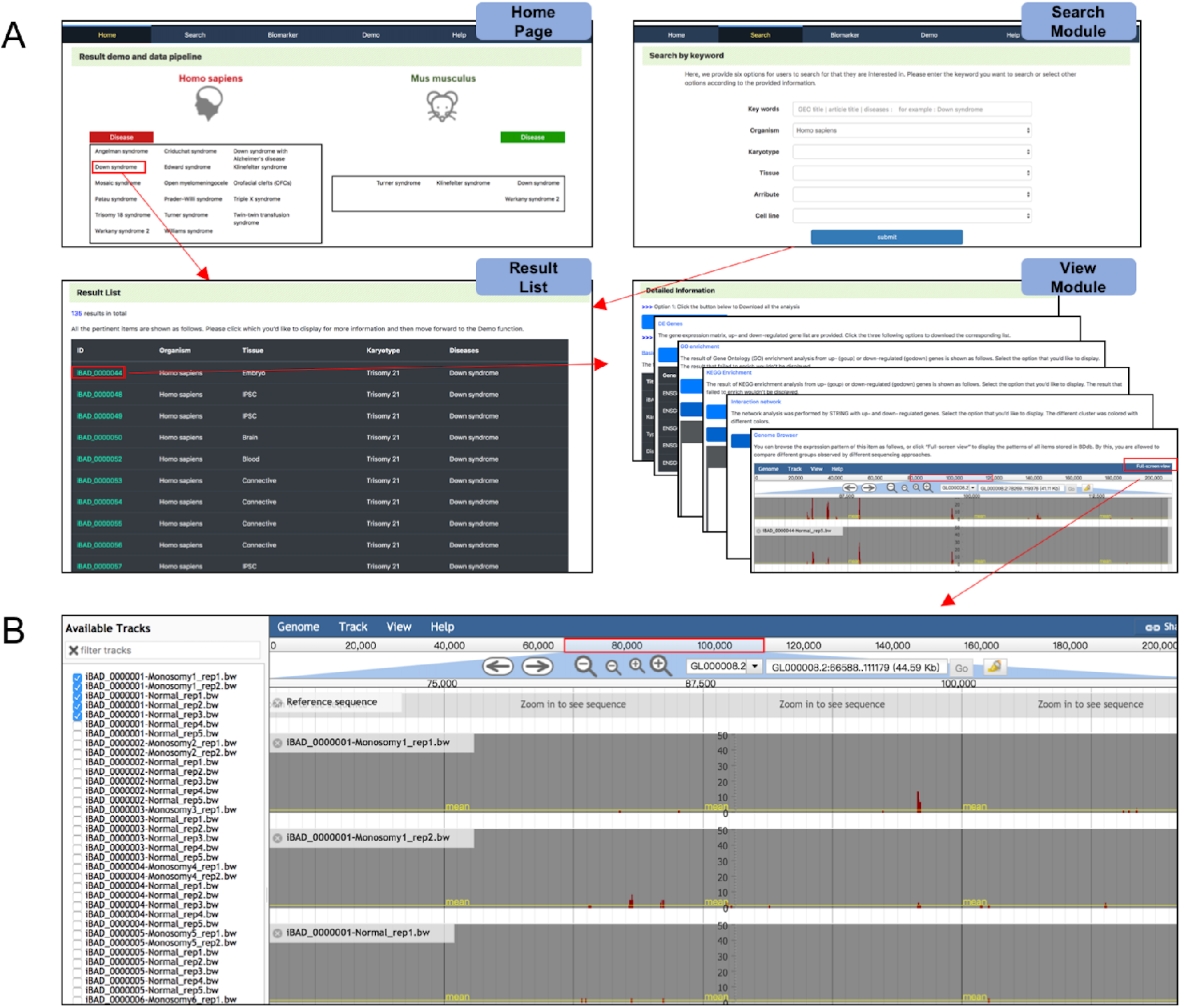
Screenshots of BDdb’s web interface. (A) An overall workflow in BDdb. Users can search the items through either home page or search module, and corresponding results would be displayed. (B) The Genome Browser interface, which enables users to interactively visualize genomic data from different studies.

### Information search

For the search module, users can search for what they are interested in by inputting keywords or by choosing the options provided. As shown in Figure 3A, users can select these options singly or multiply, including the organism, karyotype, tissue, as well as cell line. Mouse models are also classified according to karyotype as human does (Figure 3A). In terms of omics, BDdb only contains *Homo sapiens* and *Mus musculus* temporarily. However, it enables users to opt for some certain kinds of karyotypes, such as trisomy 21, trisomy 18 and monosomy X, all of which are the typical karyotypes for chromosome aneuploid. Additionally, tissues and cell lines are also the conditions used to filter searching results. After submitting the request, relevant results will be displayed.

### View module

The data from distinct sequencing types are displayed with different result modules (Figure 3A). For example, in RNA-Seq datasets, the resulting interface contains four sections: the first part, ‘Basic Information’, shows the information of karyotype, disease, organism, tissue and cell line, which are the sample’s features, as well as the GEO title, literature and searching link, which can help users to trace back to the origin of this data; the second part, ‘DE Genes’, enables users to look through or download the gene expression matrix as well as up- and down-regulated gene table; in the third section, to dive into the function/pathway enrichment, GO and KEGG enrichment are offered, with the bubble chart, bar chart and cnetplot being showed; the fourth section displays the network analysis of differentially expressed genes performed by STRING (29); and the last section, ‘Genome Browser’, can intuitively display the expression pattern with the graphical interface. Apart from the datasets from RNA-seq, we also provide fundamental analysis for other datasets. Particularly, omics datasets from different studies can be displayed in the Genome Browser as per user’s requirements, to further mine other useful information (Figure 3B).

### Birth-defect diseases biomarker mapper

The biomarkers are stored in the biomarker-mapper module. Users can search and view for markers of interest by selecting species, diseases, tissues and cell types from the pull-down menu.

Besides, users can download all analysis results and biomarkers via the “Download” bottom. BDdb also provides a detailed tutorial and common questions with answers for the usage of the database on the “Help” page.

### Case Study: exploring more biomarkers for diseases diagnosis using BDdb

In hopes of mining some useful clues of diseases leveraging BDdb database, we targeted at Down Syndrome in *Homo sapiens*, which has drawn massive attention across the world over the years. Taking the fibroblasts as an example, a total of 13 GSE samples was linked to trisomy 21, including various sequencing types such as RNA-Seq and DNase-Seq and more. We consolidated the up-regulated DE genes from 8 GSE samples performed by RNA-Seq and DNA microarray, and then sorted them by counts. The count of 21 genes, in total, was over or equal to 4 (Figure 4A). Among them, the *TTC3* (*tetratricopeptide repeat domain 3*) and *IFI27* (*interferon α-inducible protein 27*) ranked 1st with the same count at six. *TTC3* is a gene located on 21q22.2 within the DS critical region (DSCR) and plays a critical role in neural development. Besides, *TTC3* is commonly regarded as a candidate gene of DS (Down Syndrome) and AD (Alzheimer’s Disease) (30,31). Meanwhile, *IFI27* has been experimentally proved to involve the interferon response in trisomy 21 (32). We found that both *TTC3* and *IFI27* have higher expression levels in T21 compared with D21 in GSE55504, in accordance with a similar pattern of chromatin accessibility in GSE55425 assessed by DNase-Seq (Figure 4B), implying that the extra copy of chromosome 21 or other transcript regulators in DS might predominantly confer to this difference. In addition to *TTC3* and *IFI37*, other genes such as *SH3BGR* (33,34) and *APP* (35) (Figure 4A) are typical biomarkers for Down Syndrome as well. As these well-studied T21 marker genes can be captured by BDdb, the rest that has not been reported yet, such as *OLFM2* and *HAS1*, might be prospective biomarkers for trisomy 21. Taken together, we can theoretically seek out more biomarkers associated with one particular disease using BDdb.

**Figure 4.**
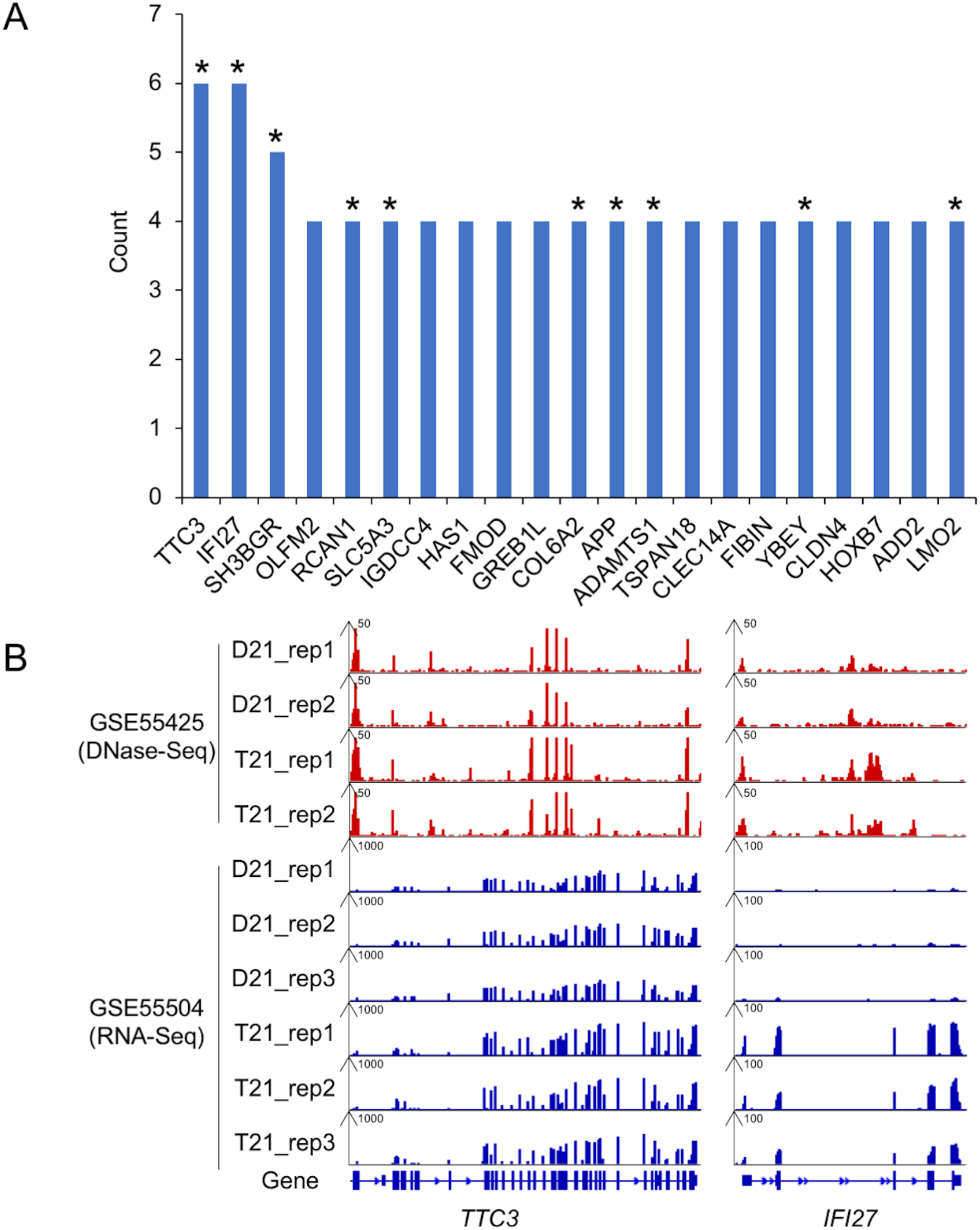
Prospective biomarkers in Down syndrome. (A) A total of 21 up-regulated genes counts more than 3 in fibroblasts from 8 GSE samples, and half of which, marked with asterisk, have been reported as prospective biomarkers for trisomy 21. (B) Gene expression patterns of *TTC3* and *IFI27* gene observed from DNase-Seq data in GSE55425 and RNA-Seq data in GSE55504.

## SUMMARY AND FUTURE PERSPECTIVE

Congenital defects, most of which are derived from chromosomal aberrations, are catastrophic for any family. Fortunately, these diseases can be effectively screened at the gestational stages via various approaches, including karyotype, chromosomal microarray analysis (CMA), next-generation sequencing (36), etc. Additionally, cell-free DNA (cfDNA) screening at the basis of maternal blood samples has been advanced to target the detection of chromosomal abnormalities in the fetus (37,38). Now such chromosome variations such as trisomies 13, 18, and 21 (39) have been validated using cfDNA testing. Moreover, a large number of data sets have been generated, leading to desperate requirements for a comprehensive database to integrate these multi-omics data. Aiming to assist clinicians and researchers, we developed the BDdb consisting of multi-omics data and datasets of cfDNA. The data related to birth defects were curated, re-analyzed and displayed in BDdb. Users can pick up the data they interested to gain a deeper understanding of the underlying molecular mechanism. Data retrieval and download are available in BDdb. Meanwhile, the list of differentially expressed genes (DEGs), the enrichment of GO terms and KEGG pathways and their interaction networks are provided. Notably, users can also utilize the genome browser to intuitively compare diverse samples as well as the data from various sequencing methods, such as RNA-Seq, ChIP-Seq and DNA methylation, or identify their correlation.

In conclusion, BDdb is a practical and instrumental tool for the research community committed to studying birth defects. To date, BDdb is, to our knowledge, the first comprehensive database related to birth defects. It will be updated constantly according to the frequency of publications associated with chromosomal aberrations. Aside from the existing data, we will also add proteomics data to expand its repository. Moreover, other species and diseases will be built in to provide more information to users. Ultimately, we hope that BDdb, serving as an auxiliary tool, could speed up the research of chromosomal aberrations to fundamentally and deeply study the congenital defect.

## ACKNOWLEDGEMENTS

We would like to thank China National GeneBank and BGI Research.

